# Mapping neurons and brain regions underlying sensorimotor decisions and sequences in *Drosophila*

**DOI:** 10.1101/215236

**Authors:** Tihana Jovanic, Jean-Baptiste Masson, James William Truman, Marta Zlatic

**Author notes:** Correspondence (T.J.), (M.Z.).

## Abstract

Nervous systems across the animal kingdom have the ability to select appropriate actions and sequences of actions in response to sensory cues. The circuit mechanisms by which nervous systems achieve choice, stability and transitions between behaviors are still incompletely understood. To identify neurons and brain areas involved in controlling these processes, we developed an approach where we combined a large-scale neuronal inactivation screen with an automated action detection of sensorimotor decisions and sequences in response to a sensory cue in *Drosophila* larva. We analyzed behaviors from 2.9×10^5^ larvae and identified 51 candidate lines for sensory processing and 24 candidate lines for competitive interactions between actions during sensorimotor decisions. We also detected phenotype categories for sequence transitions consistent with a model of sequence generation where transitions and reversals are independently controlled. These findings provide the basis for understanding how sensorimotor decisions and sequence transition are controlled by the nervous system

## 1. INTRODUCTION

In order to enable reliable and coherent responses of organisms to sensory cues, the choice of one action must be accompanied by a full suppression of all competing physically mutually exclusive actions. Often, animals respond to stimuli, not with single actions but with sequences of actions in which case, the transitions between actions need to be precisely controlled. These actions can vary between individuals and also within individuals upon repetition of the stimuli and thus have a probabilistic component. The circuit implementation of competitive interactions between neurons that promote mutually exclusive actions and the circuit mechanisms that control action stability and transitions from one action to the next are still incompletely understood.

For example, whether the choice of a behavior, its stability and transition into different behaviors are implemented in specialized decision-making centers or distributed across the nervous system form the sensory to the motor side is still a heavily debated subject (Cisek, 2007; Cisek and Kalaska, 2010; Gaudry and Kristan, 2009; Gold and Shadlen, 2007; Kristan, 2008; Lebedev and Wise, 2002; Livant, 2007; Miller et al., 1960; Reyn et al., 2014). We have previously identified a circuit for behavioral choice (between 2 actions) and sequences in response to an airpuff at the earliest stage sensory processing site which supports a model in which competitive interactions are distributed across the nervous system (Cisek and Kalaska, 2010; Jovanic et al., 2016). To examine the generalization of this computing architecture, it is necessary to identify other sites of competitive interactions and understand how the outcomes of these putative circuits for behavioral choice are integrated to give rise to a unified decision about which action to perform.

In addition, when actions are executed in sequences, how does the nervous system specify the order of the individual actions in the sequence? There have been different theoretical models of sequence generation proposed: parallel queuing (Lasley, 1951; Seeds et al., 2014), synaptic chains (Long et al., 2010) and more recently chains of disinhibitory loops (Jovanic et al., 2016), but how flexible behavioral sequences are implemented in the nervous system remains an open question.

A first step towards understanding the circuit mechanisms of action selection and probabilistic sequence generation is to identify neurons and nervous system areas involved in promoting, stabilizing and suppressing specific actions.

Here, we used the GAL4/UAS system available in the *Drosophila* to selectively and constitutively silence neurons in populations of behaving *Drosophila* larvae during behavioral responses to an air-puff stimulation using tetanus-toxin. The airpuff can evoke 4 different actions in the larva that can be organized in various sequences which makes it suitable for studying both sensorimotor decisions and sequences. We have previously used this assay to study the circuit mechanisms underlying the choice between the two most prominent actions that occur in response to air-puff and a sequence of these two behaviors (Jovanic et al., 2016). Here we expanded the study on the choice between multiple behaviors and longer behavioral sequences (of up to 4 different actions). We designed a behavioral screen using a library of *Drosophila* lines (Jenett et al., 2012; Li et al., 2014). Based on anatomical pre-screen of a collection of images of the larval stages of the GAL4 *Drosophila* lines we selected those that target small subset of neurons. 567 driver lines were screened and the behaviors of tens of thousands of animals recorded.

To quantify larval responses, we used supervised machine learning to reliably detect the actions that can be evoked by air puff and their sequences. In addition to new sensory lines, we identified 75 candidate GAL4 lines that target neurons required for behavioral responses to mechanosensory stimuli. Amongst these, we identified candidate elements of functional circuit modules for sensory processing (51 GAL4 lines) and sensorimotor decisions (24 GAL4 lines). We also identified candidate hit lines for specific sequence transitions (127 GAL4 lines total). The hit lines span different regions of the ventral nerve cord and the brain which suggests multiple sites of competition that are distributed across the nervous system.

These candidate lines provide a valuable resource for those interested in studying neural mechanisms underlying sensorimotor behaviors as it can restrict the focus for those interested in specific aspect of behavior (i.e. sensory processing, decisions, sequence generation) from the exploration of all the neurons in the nervous system to specific neurons and brain regions. Therefore this database is a prerequisite for the comprehensive circuit studies underlying sensorimotor decisions and action sequences during behavioral responses to a mechanical sensory cues as the candidate neurons can be used as starting points for circuit mapping and analysis by combining the results of silencing behavioral genetics experiments with the different methods available in *Drosophila* larva to relate structure and function of neuronal circuits: EM reconstruction, optogenetics, electrophysiology, calcium imaging, immunohistochemistry and modeling (Heckscher et al., 2015; Jovanic et al., 2016; Ohyama et al., 2015; Zwart et al., 2016).

## 2. RESULTS

### 2.1 An assay for sensorimotor decisions and sequences, using machine-learning based characterization of the behavioral response to an air-puff

In order to functionally map neurons specifically to the different aspects of sensorimotor behaviors, we presented stimuli to populations of freely behaving wild type larvae and larvae with inactivated neurons and monitored their behavioral responses (Figure 1 A-B)

**Figure 1.**
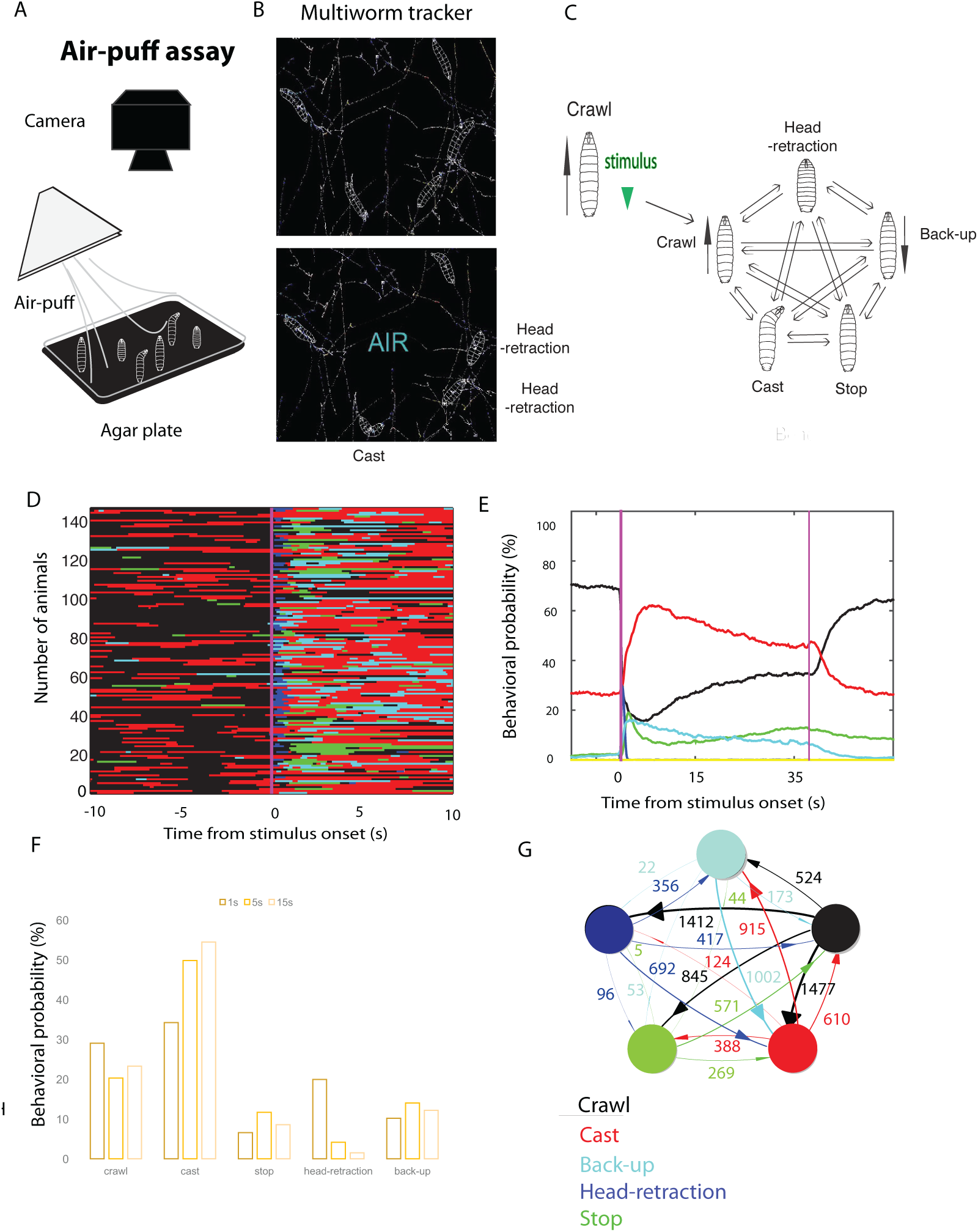
Characterisation of the behavioral response of wild-type larvae to an air-puff. **A.** Behavioral set-up **B**. Still from a movie of contours of larvae acquired with the Multiworm tracker software **C**. In response to air-puff larvae perform a probabilistic sequence of four actions **D**. Ethogram of the behavioral response to an air-puff (10s after stimulus onset). Here a subset of animals tested is shown for clarity (N larvae shown is 145). Different colors represent different actions: Blue-Head-retraction, Red-Cast B, Cyan-Back-up, Black-Crawl and in Green-Stop **E**. Behavioral probability (mean) before, during and after stimulation **F**. Behavioral probabilities in the first second, first 5 seconds and first 15 seconds after stimulus onset **G**. Transitions probabilities between five actions: Crawl to the Head-retraction 35.9 %, Crawl to Cast 31.4 %, from Head-retraction to Cast 42 % and Head-retraction to Crawl 27.1 %, to back-up 25.3% and to stop 5.6 %. The cast has the highest probability of transitioning into a back-up (45.8%), a moderate probability of transitioning into a crawl (29.7 %), a lower probability of transitioning into a stop (18.8 %) and a lowest probability of transitioning into a head-retraction (5.6%). The Stop has a high likelihood of transitioning into a cast and lower probability of transitioning into a crawl. From back-up there is the highest probability to transition into a cast (79.9 % than into a crawl (13.6%) and with lower probability into a Stop and Head-retractions (4.5 % and 2%). The colors are as in D and E. The plot represents all the transitions in the first 3 s after stimulus onset

We used a supervised learning algorithm that relies on a limited set of features, and is able to handle low resolution images of larva to specifically detect 5 action categories that we observed in response to air-puff: head-retraction, stop, cast, back-up and forward crawl. These different discrete actions emerge naturally as separate clusters in a simple embedding space derived from few features. The actions are mutually exclusive and non-overlapping (Supplementary figure 1 A).

We have previously studied and analyzed the immediate response of larvae to the air-puff using a length variation method (Jovanic et al., 2016; Ohyama et al., 2013) (see Materials and methods for the link between new action definitions and previous ones). Here we analyzed a dataset of wild type larvae (control: w;;attP2-TNT larvae, (progeny of male w;;attP2 crossed to female UAS-TNT-e flies (13 831 larvae), using the newly developed method and quantified previously undetected actions: stops and back-ups, in addition to casts, head-retractions and forward crawls (Fig 1 C-E,G). Each of these actions occurs at different probabilities and these probabilities vary over the time of the response (Figure 1 D-F). For example, head-retractions have a higher likelihood of occurring early in the response while the stop occurs with a high probability early, its probability than drops and increase in the second half of the response (Figure 1 E).

We computed behavioral probabilities for each of the five actions during first second, during five and during fifteen seconds after stimulation (Figure 1F) and because the head-retractions occurs primarily immediately after stimulus onset we focused the behavioral probability analysis of the screen data on the shortest time window.

We further investigated the effect on air-puff intensity on the behavior. While the probabilities of some actions vary significantly with stimulus intensity (i.e. higher head-retraction and lower stopping at high intensity of air-puff), the amplitude of actions doesn’t seem to change with stimulus intensity (Supplementary figure 1 B)

We computed transition probabilities between the different actions during the first 3 seconds after stimulus onset and found that from the crawl the larvae will be transitioning strongly into a cast or a head-retraction (35.9 % and 31.4 %) respectively, less strongly into a stop and with a lowest probability into a back up (Figure 3). Head-retraction can transition into each of the four other actions with the highest likelihood of transitioning into a cast (42 %) and lower probability of transitioning into the baseline crawl and back-up (27.1 % and 25.3 %) and a low transition probability to stop 5.6 %). The cast has the highest probability of transitioning into a back-up (45.8%), a moderate probability of transitioning into a crawl (29.7 %), a lower probability of transitioning into a stop (18.8%) and a lowest probability of transitioning into a head-retraction (5.6 %). The stop has a high likelihood of transitioning into a cast and lower probability of transitioning into a crawl. From back-up, the highest probability is to transition into a cast (79.9% than into a crawl (13.6 %) and with lower probability into a stop and head-retractions (4.5 % and 2%). This suggests that the first action in a sequence is most frequently a head-retraction or a cast (Jovanic et al., 2016) and that if a head-retraction is the first action it will be followed by the cast with the highest probability. If cast is the first action in the sequence than there is higher likelihood that the second action will be a back-up (or stop) than a head-retraction. Because transition probabilities between cast and back up are high in both directions, it suggests that there are multiple transition events between cast and back-up in one sequence with higher probabilities of transitioning from the back-up onto the cast. The transitions between head-retractions and cast, head-retractions and back-ups and back-ups and casts are asymmetrical, meaning that the transitions are more likely in one direction that the other (head-retractions to cast and back-up and back-up to cast).

We further investigated the types of sequence motifs produced in response to an air-puff. We looked at the first four actions in the sequence (after stimulus onset) and identified the 10 most common four action sequence motifs that occur in the w;;attP2-TNT larvae upon presentation of the stimulus composed of head-retractions, casts, back-ups, stops and the baseline behavior crawl. The most frequent motif is the head-retraction — cast - back-up-cast (Supplementary table 4).

In summary, in response to an airpuff, the larvae perform a probabilistic sequence that can be composed of up to four different actions where the head-retraction is most likely to occur immediately after stimulus onset while the cast, back-up and stop can occur later in the response. The behavioral response to an air-puff is thus well suited to use as an assay to study sensorimotor decisions between multiple actions and generations of longer behavioral sequences.

### 2.2. Inactivation Screen for mapping neurons and brain regions that underlie the behavioral response to air-puff

In order to identify neurons specifically involved in sensorimotor decisions and sequences we applied the automated algorithm for behavioral detection to a behavioral inactivation screen dataset of larval responses to an air-puff stimulus. We analyzed behavioral responses of larvae in which we silenced small subsets of neurons and individual neurons. We tested 567 GAL4 lines that we selected for sparseness and expression quality based on an anatomical pre-screen out of the collection of the JRC GAL4 lines (Li et al., 2014). Expression patterns of the GAL4 lines used in this study are characterized in details and neuroanatomical information are available on http://wwwjanelia.org/gal4-gen1.

Ethogram of the full behavioral response to the air-puff of all the lines in the screen (Supplementary figure 2A, left) shows that the dominant behavior immediately after the stimulus onset is head-retraction and the second dominant behavior is a cast. The head-retraction is most likely to be followed by a stop or a cast, while the second dominant behavior at the second position would be the back-up and crawl (stop if the first dominant in the second position is a cast (Supplementary figure 2 A, right). Head-retractions, stops and back-up occur only after stimulus. Crawl and cast also occur prior to the stimulus onset, as these actions are part of larval foraging behavior.

**Figure 2.**
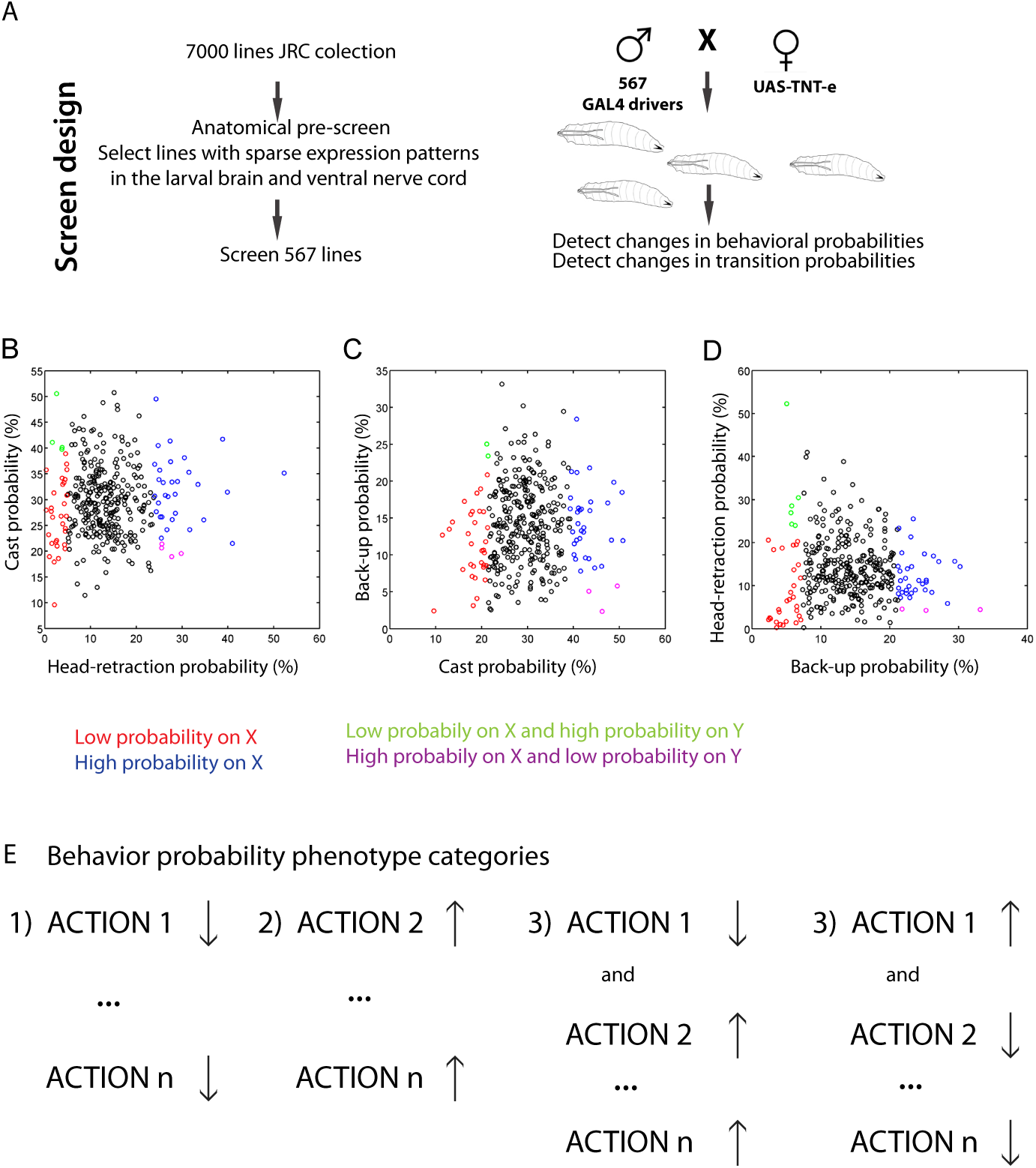
**A**. Experimental strategy and screen design **B**. Scatterplot of Head-retraction probability against Cast probability. The 10% of lines with lowest and highest probabilities of head-retraction are shown in red (+green) and blue (+magenta) respectively. The lines with low head-retraction and high cast probabilities are shown in green. The lines with high head-retraction and low cast probabilities are shown in magenta **C**. Scatterplot of cast probabilities against back-up probability. The 10% of lines with lowest and highest probabilities of casts are shown in red (+green) and blue (+magenta) respectively. The lines with low cast and high back-up probabilities are shown in green. The lines with high cast and low back-up probability are shown in magenta **D**. Scatterplot of Back-up probabilities against head-retraction probability. The 10% of lines with lowest and highest probabilities of back-up are shown in red (+green) and blue (+magenta) respectively. The lines with low back-up and high head-retraction probabilities are shown in green. The lines with high back-up and low head-retraction probabilities are shown in magenta **E**. Behavioral probability phenotype categories: 1) less response phenotype: lower probabilities in one or more actions 2) more response: higher probabilities in one or more actions 3) competitive interaction phenotype: lower probability of one action and higher probability of at least one action or vice versa

Some of the tested lines did not have the same hierarchy of behavior as the majority including the control prior to stimulus onset. This is most likely due to locomotor defects (Supplementary Spreadsheet_locomotor defect lines). We report these lines as these neurons are good candidates for examining circuitry underlying locomotion. For the purpose of studying stimulus evoked decisions and sequences, we excluded those lines from further analysis as their defects would interfere with animal’s capacity to respond to stimulation.

We determined families of hits by comparing the probabilities of performing each of the actions that occur in response to air-puff between test lines and the control. We considered as candidate for strong hits the 10% of lines that had the lowest or highest behavioral probabilities in head-retractions, casting or backing-up and considered as hits only those that were significantly different from the control (see Material and methods for details)

We then examined the different types of phenotypes (Figure 2 B-D, Supplementary table 2, Supplementary spreadsheet_Behavioral Probabilities) and found group of hits where: 1) one or more actions decreased, 2) one or more actions increased compared to the control and 3) one action was increased while (at least) another was decreased compared to the control. We consider the latter category of hits “decision hits” (or “competitive interaction hits”) as the phenotypes where the probability of one behavior is increased while that of others is decreased is consistent with the same neurons promoting one actions while inhibiting others. These lines are the candidate hits for studying the circuit mechanisms underlying competitive interactions between different (mutually exclusive) behaviors (see section 2.5).

In addition to the behavioral probabilities, other features of behavioral responses, for example the amplitude of individual actions could also be affected. We quantified the amplitudes of head-retractions and casts (Supplementary figure 2. B-C). The amplitude of casting was on average stronger in lines with a low probability of casting. This could be due to the fact that it takes the animal longer to recover from the action and come back to neutral position. There were some exceptions. For example, the R15F02 that has both higher amplitude and probability of casting. ON the other hand, most of lines that head-retract little also do weaker head-retractions and those that do more head-retractions do that more strongly. An exception to this is the R11A07 that has a high head-retraction probability, but relatively low amplitude. Further investigating these outliers will help understand how the neurons control both the strength and probability of the response.

Because larvae usually crawl forward prior to the stimulation (it is a baseline behavior) we excluded the crawl from behavioral probability hit detection. Similarly, we didn’t include the stopping probability in the hit detection as phenotypes in stopping probability cannot be interpreted unambiguously. At lower intensity of stimulation larvae stop more (Supplementary figure 1 B) and therefore an increase in stopping (and decrease in one or more other actions) might reflect impaired sensing of air-puff. Indeed, this type of phenotypes can be observed when silencing chordotonals-key sensory neurons for sensing air-puff (Supplementary table 1, see section 2.3). For all the hits, both the stopping and crawling probabilities are shown in Supplementary table 1 and Supplementary table 2 along with head-retraction, cast and back-up probabilities used for hit-detection.

### 2.3. At least 2 different types of sensory neurons mediate the air-puff response

We have previously identified chordotonal sensory neurons as sensory neurons that mediate the air-puff induced responses as silencing of these neurons resulted in severely impaired responses (Jovanic et al., 2016; Ohyama et al., 2013). While the response is severely impaired it is not completely abolished suggesting that some other sensory neurons could be involved in sensing air-current in addition to the chordotonals. In the *Drosophila* larva, there are multiple types of somatosensory neurons. Somatosensory neurons of type I, which are the monodendritic and are the chordotonal and the external sensory neurons and multidendritic somatosensory neurons which can be subdivided into 4 classes: I, II, III, IV depending on the level of their arborizations and the position on the body wall (Bodmer and Jan, 1987; Grueber et al., 2007; Merritt and Whitington, 1995).

Out of the lines screened we have identified 27 candidate hit lines that drive either only in sensory neurons (one or multiple types) or in sensory and some other neuron types (Figure 3A, Supplementary table 1). We found that chordotonal sensory neurons are present in at least 6 GAL4 lines (Supplementary table 1) including the positive control lines, R61D08 and iav-GAL4 lines that were described previously (Jovanic et al., 2016; Ohyama et al., 2013). Silencing of chordotonal neurons resulted in lower probability of head-retractions and casting and in some lines also of backing-up. Differences between the lines could be due to the different strength of the Gal4 drivers and to the different subtypes of chordotonal neuron targeted by the drivers (Merritt and Whitington, 1995). Multidendritic (md) and external sensory (es) neurons appear in some of the lines with perturbed response to the air-puff (i.e. R30H09, R54A01) and thus could be involved in air-puff sensing as well. While for the es neurons, we were not able to obtain a cleaner line, we obtained a line labelling multidendritic class III (md III) neurons and tested in the air-puff assay. We found that silencing of multidendritic neurons resulted in significantly less head-retractions (although less pronounced phenotype than for chordotonals) and backing-up, while the casting response remained comparable to that in the control (Figure 3, Supplementary table 1) suggesting that head-retraction and back-up are mediated also by the md III circuitry, while casts may be not. Md III have been shown to respond to light touch that induces similar response types: head-retractions, back-up, cast, and turns (away from the stimulus) (Tsubouchi et al., 2012; Yan et al., 2013). When silencing chordotonal and multidendritic class III neurons expressing tetanus-toxin using the NompC (a TRP channel that confers light touch sensitivity (Yan et al., 2013)) GAL4 driver that drives in both subsets of chordotonals and md III neurons the larvae showed an impaired response with less head-retractions, and casts, a phenotype consistent with a strong impairment in sensing air-current.

**Figure 3.**
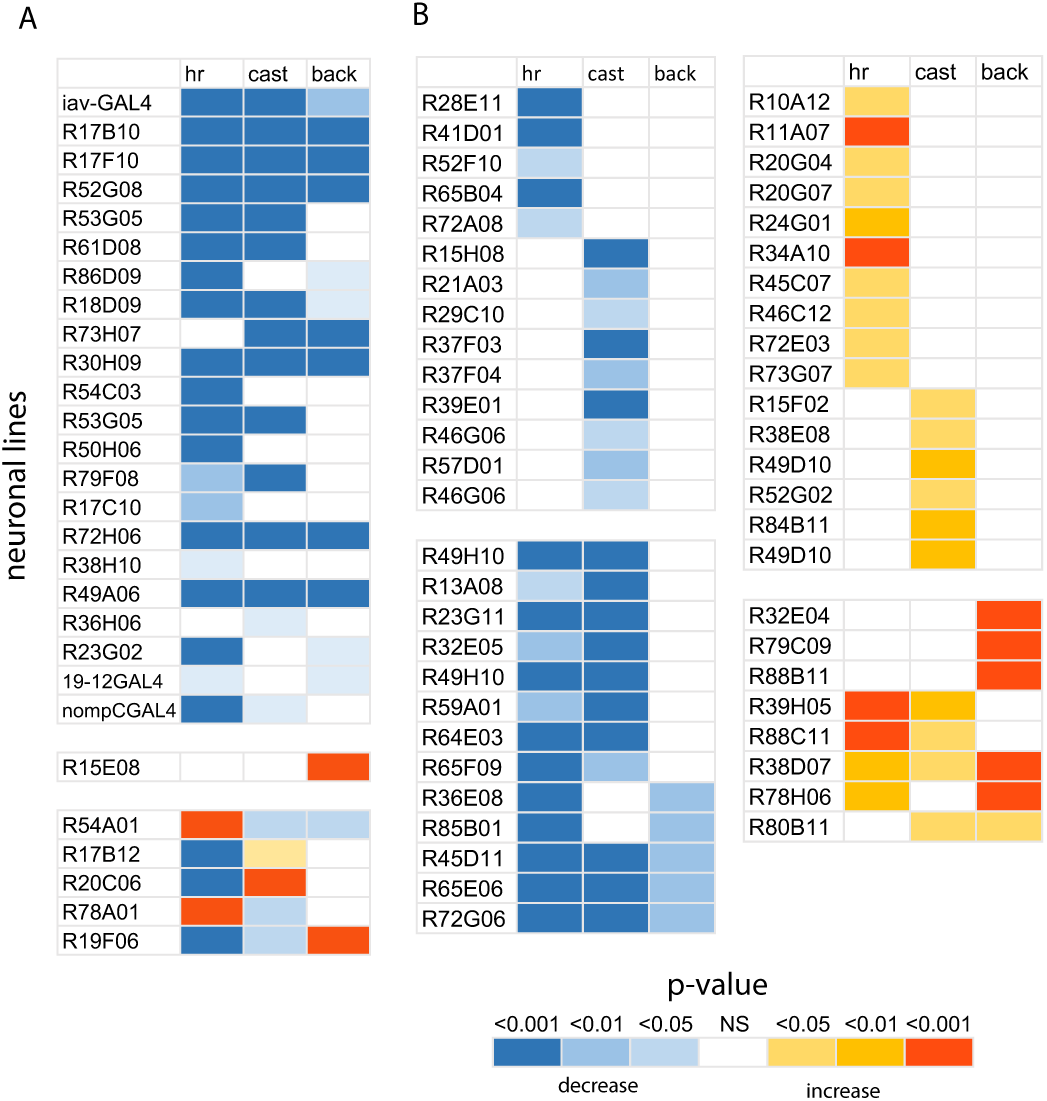
Phenotypic summaries using selected behaviors (head-retraction, cast and back-up) for the hits from the behavioral screen. The colors in the heatmap represent the p-values for each behavior for the comparison between each line shown to the right of each plot to the w;;attP2-TNT control **A**. 27 sensory lines. Known types of sensory neurons are shown in Supplementary table 1 **B**. CNS lines (51 lines). Left: Hits with lower probabilities in at least one behavior compared to the control (27 lines). Right: Hits with higher probabilities in at least one behavior compared to the control (24 lines). The behavioral probabilities can be found in Supplementary tables 1 and 2

In summary, silencing of air-current sensing neurons resulted in lower probabilities of head-retractions and for some lines also in casting and/or backing–up. When silencing the chordotonal neurons, the probability of stopping was increased in several GAL4 lines (Supplemenary table 1), which suggested that the stopping response could be mediated by different sensory neurons and in absence of chordotonal sensory neurons this type of response becomes the primary response type. Alternatively, based on the finding that lower intensities of air-puff trigger more stopping than higher, in absence of chordotonals, the larvae could stop more because they sense the stimulus less.

### 2.4. Identification of central neurons involved in the air-puff response

Out of the lines driving in interneurons in the ventral nerve cord (VNC) and the brain, we observed three main categories of hits that have significantly different behavioral probabilities from the control (Figure 2B-E). The first categories are lines in which silencing of neurons results in lower probabilities of at least one action compared to the control and the second are the lines in which silencing of neurons results in higher probabilities of at least one action. In both of these categories the other actions remain unaffected. In a third category, silencing of neurons, results in higher probability of one action and lower probability of at least one action or vice versa, suggesting that these neurons implement competitive interactions between the different behaviors.

If a neuron is required for sensory processing, the silencing of this neuron will result in a perturbed response where the larvae will react either more or less compared to a reference and the probabilities of performing an action could be affected across the behavioral repertoire (in one or more actions). Alternatively, a phenotype with higher or lower probabilities of behaviors could also be consistent with the modulation of behavioral response depending on the context or with neurons that control aspects of motor responses that will affect the speed, duration etc. of execution and affect behavioral probabilities indirectly.

Out of the hit candidate lines from the primary analysis, we identified the strong candidates in these categories that are significantly different than the control. We identified 51 such hits where the probability of at least one action was a) decreased or b) increased (Figure 3 B, Supplementary table 2).

Some of these lines have sparse expression patterns. We identified lines with more head-retraction (i.e. R11A07), more casting (i.e. R15F02) or more backing-up (R32E04). Silencing of neurons in R65B04 and R21A03 lines resulted in less head-retractions and casting respectively (Figure 3B). We didn’t identify lines that back-up less, unless the phenotype was accompanied with less of head-retractions and/or casts.

### 2.5. Identification of neurons and brain regions involved in sensorimotor decisions

We determined that the detected actions in response to air-puff are mutually exclusive (Supplementary figure 1A) and therefore we sought to identify neurons that could implement competitive interactions between these actions. We reasoned that if a neuron is required for sensorimotor decisions between mutually exclusive behaviors, inactivating that neuron might increase the probability of one action at the expense of another or several other actions.

We identified 24 hits in which larva perform less of at least one action and more of at least another action (Figure 4A, Supplementary table 2). This phenotype could be observed with different combinations of actions: head-retractions, casts and back-ups. Interestingly we never observed a decrease in back-up probabilities and increase in cast probability.

**Figure 4.**
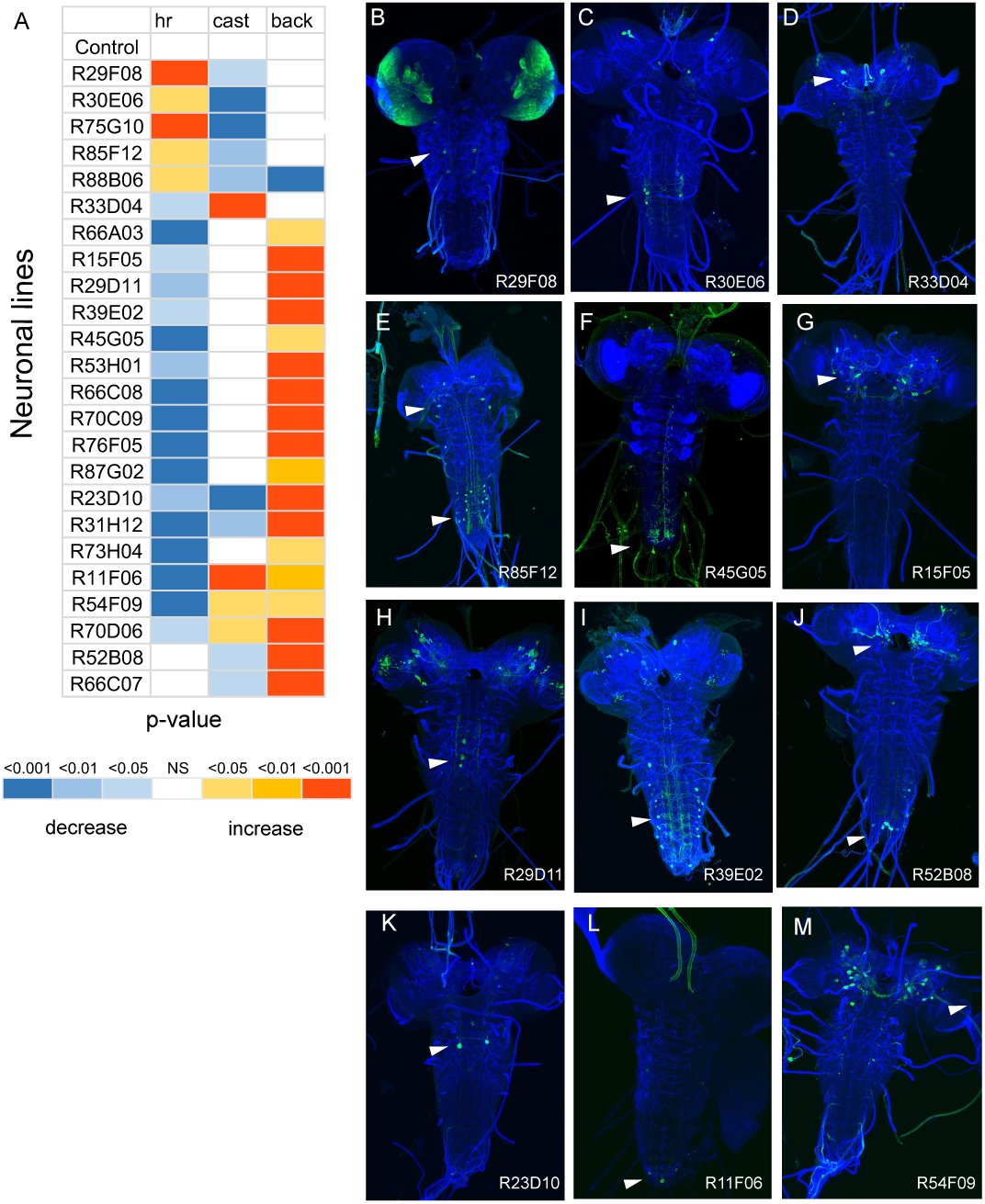
**A**. Phenotypic summaries using selected behaviors (head-retraction, cast and back-up) for the “decision” hits from the behavioral screen. The colors in the heatmap represent the p-values for each behavior for the comparison between each line shown to the right of the plot to the w;; attP2-TNT control. All the hits that have lower probabilities in at least one action and higher probabilities in at least one other action (24 lines) are shown **B-M**. Example of candidate hits with sparse expression patterns. **B-E** Hits where the competitive interactions between head-retractions and casts are perturbed **F-I**. Hits where the competitive interactions between head-retractions and backups are perturbed **K-M.** Hits where the competitive interactions between all three actions are perturbed. **B-M**. Green-GFP; Blue-N-cadherin

Twelve out of the 24 lines have sparse neuronal expression patterns (Figure 4B). For example, silencing of the neurons in R29F08 and R33D04 sparse lines results in more head-retraction and less casting and less head-retraction and more casting respectively (Figure 4).

These neurons are located in different areas of the nervous system (Supplementary table 2, Figure 4B-M), which in line with distributed models of implementation of competitive interactions and decision-making (Cisek, 2007; Cisek and Kalaska, 2010; Jovanic et al., 2016)

### 2.6. Identification of neurons and brain regions underlying sequence transitions

We computed transition probabilities between each pair of individual actions in all the different genotypes (Supplementary table 3. transition probabilities). As expected, in some of the candidate “decision hit” lines with affected behavioral probabilities (R11F06, R23D10, R29D11, R87G02, R88B06). the transition probabilities were also perturbed (Supplementary table 3, Supplementary spreadsheet_TransitionProbabilities) and these 2 statistics are intricately linked.

We have previously proposed that lateral disinhibition promotes transitions between behaviors, while feedback disinhibition prevents reversal from one element in the sequence back onto the previous one (Jovanic et al., 2016) in a probabilistic sequence of two actions. We speculated that chains of such disinhibitory loops could be a general mechanism for generating longer behavioral sequences. We therefore sought to investigate whether for other pairs of actions: 1) transitions and reversals are separately controlled and 2) whether the maintenance of an action (through a positive feedback) allows the proper progression through the sequence and prevents reversals.

Consistent with our hypothesis that transitions and reversals are controlled independently, we identified neuronal lines where silencing of neurons affected only transitions or only reversals from one element in the sequence onto another (Supplementary table 3, in red transition hits are shown, in green reversal hits, Supplementary spreadsheet_TransitionProbabilities). We first analyzed hits with affected transition probabilities from the head-retraction to the cast and to the back-up (Figure 5 D, Supplementray table 3). We identified 62 such candidate hits (24 that affect head-retraction to cast transitions and 38 that affect head-retraction to back-up transitions). Interestingly while the head-retraction to cast hits had lower transition probabilities compared to the control, the majority of head-retraction to back hits had higher transition probability compared to the control (except for 5 lines). This suggests that the transitions from head-retraction to back-up are strongly inhibited in the wild-type larvae during early response to an air-puff.

**Figure 5.**
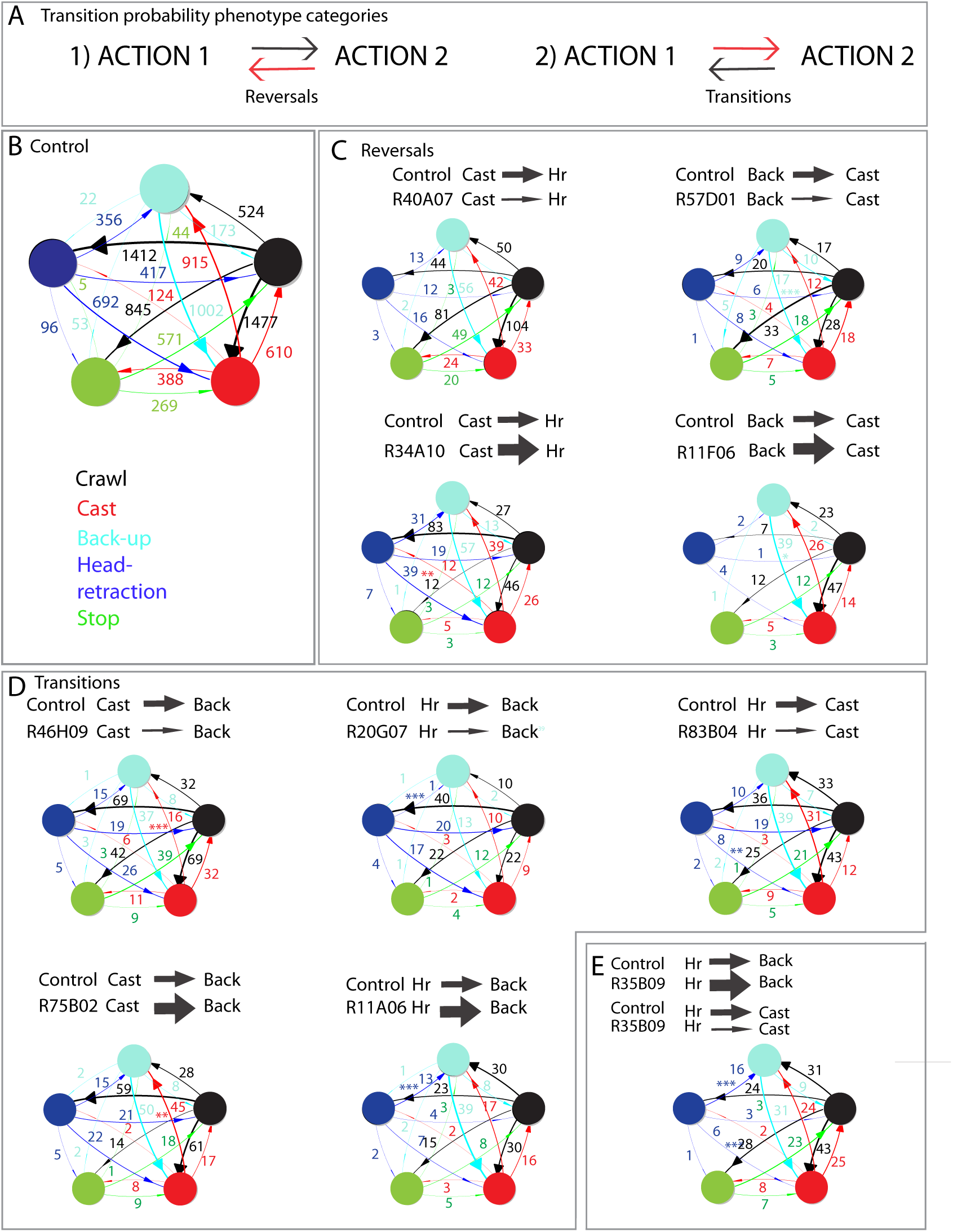
**A**. Transition probability phenotype categories shown: 1) Only reversal affected, 2) Only transitions affected **B**. Transition probabilities for the Control: w;; attP2-TNT control **C.** Reversal hits: examples for Cast to Head-retraction and Back-up to Cast, **D**. Transition hits: Examples for Head-retraction to Back-up, Head-retraction to Cast and Cast to Back-up **E**. Example of a hit with an increase in transitions from head retractions to back-up and decrease in Head-retraction to Cast transitions

We then analyzed candidate hit lines where both head-retraction to cast and head-retraction to back-up were affected to see whether there were competitive interactions that underlie the choice of the action to transition to, from the head-retractions. We found that the lines where both head-retraction to cast and head-retraction to back-up transitions were affected, had one type of transition increased and the other decreased compared to the control consistent with the existence of competition between the transitions into a back-up or a cast. Out of the 18 lines, only one had an increase in the head-retraction to cast transitions and a decrease in head-retraction to back-up transitions. The remaining 17 lines showed an increase in head-retraction to back-up transition and decrease in head-retraction to cast transitions, phenotypes consistent with most of the lines promoting transitions from head-retraction to cast and inhibiting transition to back-up (Supplementary table 3, Figure 5 E).

Interestingly, in behavioral probability hits we never observed a decrease in back-up at the expense of cast or head-retraction (except a decrease in both back-up and cast and increase in head-retraction) (Figure 4A), which suggests that, at least in the early response to air puff the head-retraction and cast are dominant behaviors and that back-up response gets activated by a head-retraction (25.3%) or a cast (42%) possibly through disinhibition like we have shown for the two action sequence (Jovanic et al., 2016).

We then analyzed hits with affected transition probabilities from back-up to head-retractions and cast to head-retraction (reversals, these transitions occur rarely in the wildtype: 2 % and 5.6% respectively) and found 6 lines that affected only transitions from cast to head-retraction out of which 2 had increased transition probabilities consistent with their role in preventing reversals (Figure 5C, Supplementary table 3 sequence transition hits). We didn’t identify any candidate hits that affected only the back-up to head-retraction transitions.

We next analyzed transitions between cast and back-up. The cast is more likely to occur after the first action in the sequence (the head-retraction) than the back-up (Figure 1. G). Unlike the reversal to head-retraction the reversal to cast occurs very frequently (in 79.8% of cases while the transition probability from cast to back-up is 45.8%). Out of 14 candidate lines that had only the back-up to cast transition probabilities affected (reversals) 1 had increased transitions (Figure 5C, Supplementary table 3.). We identified 27 candidate lines in which the cast to back-up transitions were affected, 16 of which show a decrease (promote transitions) and 11 show increased transitions from cast to back-up.

The fact that we find phenotypes where only one transition or reversal is affected between a pair of actions suggests that these processes are controlled separately.

To investigate how the changes in transition probabilities between head-retractions, casts and back-ups as well as behavioral probabilities of the individual actions, affect the type of sequence produced, we examined across all the lines screened, the probability of performing three most frequent types of four action sequence motifs composed of head-retractions, back-ups and casts (Supplementary figure 4). We identified 61 lines where the probability of performing the head-retraction – cast - back-up - cast, head-retraction -back-up - cast - back-up and/or cast-back-up-cast-back-up motifs is either increased or decreased compared to the control (Supplementary figure 3). Out of the 61 lines, 34 lines had also higher or lower behavioral and/or transition probabilities compared to the control and thus the changes in the sequence motifs probabilities result from the changes in probabilities of executing one or more actions in response to the stimulus and/or transition probabilities between actions. The remaining 27 lines had no phenotype in the observed head-retraction, back-up and cast probabilities as well as the transitions probabilities between pairs of these actions, suggesting that the neurons in these lines contribute the sequence generation by different mechanisms that do not significantly affect the behavioral probability of individual actions or the transition probabilities between the three actions and that are yet to be elucidated Thus the candidate lines identified in Supplementary table 3 represent an additional resource mapping neural circuits underlying sequences and for understanding the neural mechanisms of sequence generation.

## 3. DISCUSSION

In order to identify neurons and brain regions underlying sensorimotor decisions and probabilistic action sequences we performed a high-throughput inactivation screen where we silenced individual neurons and groups of neurons (using tetanus-toxin) in 567 genetic GAL4 lines in *Drosophila* larva and looked at the effects of these manipulations on decisions and sequences occurring in response to a mechanosensory cue.

We characterized the behavioral response of wild-type larvae to the stimulus (air-puff) and found that larvae perform a probabilistic sequence of four different actions. We developed and used an automated approach that detects and distinguishes 5 different discrete behaviors that larvae perform in response to the air-puff (the four air-puff induced actions and the baseline behavior). Evidences suggest that the discrete action description is relevant when compared to a continuum approach as parameters associated to larva dynamics tend to naturally cluster. We used this analysis to describe phenotypes that result from manipulation of different populations of neurons or single neurons. We find phenotypes that are consistent with a specific role of neurons in sensory processing, sensorimotor decisions (competitive interactions) and sequence transitions. Neuronal expression data for all of the GAL4 lines used in this screen have been previously published (http://www.janelia.org/gal4-gen1). The number of neurons that were targeted in our tested lines varies from 2 to 7 pairs on average and smaller number of the GAL4 lines the driver is restricted to a single neuron type.

Our screen therefore identified candidate regulatory and control centers in the nervous system for sensorimotor decisions and probabilistic sequences. These candidate neurons and nervous system regions can restrict the focus for those interested in studying specific aspects of behavior: sensory-processing, sensorimotor decisions and sequences from the entire nervous system to a relatively small number of elements. For example, connectivity and activity mapping efforts can be focused just to the relevant subset of neurons.

We developed a framework for identifying circuit elements underlying decisions and sequence transitions. Sensory-processing, sensorimotor decisions and sequence generation are intertwined processes as the latter two will depend on how the sensory information is processed, and the sequence production mechanistically might depend on competitive interaction between distinct actions (competitive queuing, chain of disinhibitory loops (Jovanic et al., 2016; Lasley, 1951; Rosenbaum et al., 2007; Seeds et al., 2014; Bullock, 2004). Nevertheless, we used the reasoning below to identify neurons selectively involved in competitive interactions that underlie decision-making and sequence generation.

We reasoned that, if the stimulus cannot be processed and thus perceived accurately the animals might respond less, by performing less of all or some of the actions. If the sensory processing is affected in the opposite way (hypersensitivity), animals might respond more, and perform more of all or some of the actions normally observed. Thus, the neurons that gave such inactivation phenotypes (of less of one or more actions; or more of one or more actions) were assigned a candidate role in sensory processing.

On the other hand, inactivation of neurons involved in mediating competition between actions is expected to result in increased probability of one action and decreased probability of one or more other actions (or the converse).

We identified 24 hits (GAL4 lines) that affected competitive interactions that underlie sensorimotor decisions. These GAL4 lines drive in neurons that are located in the ventral nerve cord (both abdomen and thorax region), brain, suboesophagial ganglion. This suggests that the networks for sensorimotor decisions are distributed across the nervous system. In a previous study, we mapped a circuit for behavioral choice and sequence generation located in the early processing centers (Jovanic et al., 2016) suggesting that the choice between two types of responses is not centralized in higher-order centers but distributed in the nervous system. It is still unknown how the circuitry early at the sensory processing site interacts with other nervous system areas to produce coherent decisions. A perceptual decision could be made about the stimulus, categorizing it into a high-risk, very aversive, which would result in an escape type of response or a low risk or low aversive which would result into a startle type of response (stop and head-retraction). Alternatively, the decision about which behavior to perform could be modulated at the early sensory processing site by top-down pathways and no other site of competitive interactions would exist that would contribute to this type of decision to a somatosensory stimulus. Finally, competitive interactions between different sensorimotor pathways could be distributed across the nervous system (Cisek and Kalaska, 2010) and other sites in the brain, thorax or towards the motor side would exist that would contribute to the decisions. In this case the competitive interactions at the early sensory processing site would contribute to the decision-making process that is the result of the integration of competitive interactions across the nervous system, including other areas i.e. brain, thorax and pre-motor areas. The identification of candidate neurons that could mediate competitive interactions in other regions of the nervous system is in line with a distributed decision-making process and suggest that other sites of competitive interactions (in other regions of the nervous system) contribute to the decision of what to do in response to mechanosensory stimulus. The identification of candidate elements provides the groundwork for understanding how the decision-making process is organized in the nervous system.

The idea that decisions are made “through a distributed consensus that emerges in competitive populations” and that interactive behaviors require sensorimotor and selection system to function in parallel (Cisek and Kalaska, 2010) have emerged in various fields (Briggman et al., 2005; Cisek, 2007; Gaudry and Kristan, 2009; Ledberg et al., 2007; Pezzulo and Castelfranchi, 2009; Reyn et al., 2014), but it has been challenging to elucidate the neuronal architecture that would implement such decisions. The *Drosophila* larva, because of its numerical simplicity, small size and the existence of multiple experimental approaches for structural and functional connectivity studies, behavioral genetics, optogenetics etc. is an ideal system for investigating how the outcomes of these competitive interactions at the different sites are integrated across the nervous system to give rise to coherent behavioral choices. This study provides the towpath for analyzing the functional neural architecture underlying decision-making.

The neural architecture that controls the productions of probabilistic action sequences and establishes the order of the individual elements in the sequence is also poorly understood. In our previous work on a two element sequence in response to an air-puff, we have proposed that transitions to the next element in the sequence and reversal to the previous element are controlled independently through two different motifs: lateral disinhibition from the neuron driving one behavior onto the neuron driving the following behavior and feedback disinhibition that provides a positive feedback that stabilizes the second behavior and prevents reversals back onto the previous actions. Based on this, in the screen we specifically looked at candidate lines with phenotypes where either only transitions or reversals between two actions were affected to test whether this may be a general mechanism involved in transitions between multiple actions. We have identified categories of phenotypes that inform us about the implementation of sequence transitions in the nervous system. We looked at transitions from the head-retraction to cast and back-up, and transitions from cast to back-up and the corresponding reversal transitions. We found phenotypes consistent with neurons that prevent reversals (from cast to head-retraction and back-up to cast). In addition, we found phenotypes that suggest that asymmetric competitive interactions exist between transitions to cast and back-up (from head-retractions) where the cast inhibits the back-up but the back-up doesn’t inhibit the cast.

In summary, our screen provides a starting point for identifying the mechanisms underlying the competitive interactions between behaviors as well as the transition between individual actions in probabilistic sequences. While the number of neurons that were targeted in our tested lines varies from 2 to 7 pairs on average, in the case when the lines label multiple neuron types, intersectional strategies can be used to further refine the expression patterns. In the larva, a volume of electron microscope data has been acquired and more than 60% of the nervous system has been reconstructed through collaborative efforts (Berck et al., 2016; Eichler et al., 2017; Fushiki et al., 2016; Heckscher et al., 2015; Jovanic et al., 2016; Ohyama et al., 2015; Schlegel et al., 2016; Schneider-Mizell et al., 2016; Zwart et al., 2016). The identified candidate neurons can therefore be reconstructed in the EM and the synaptic patterns of interactions between neurons involved in competitive interactions can be mapped. Combined with EM reconstruction, electrophysiology, and modeling the candidate lists of neurons can be used to relate structure and function and unravel the principles of how the nervous system makes decisions and produces behavioral sequences across the nervous system.

## 4. METHOD DETAILS

### Drosophila Stocks

Fly stocks We used 567 GAL4 from the Rubin collection lines listed in Supplementary Dataset 2 (available from Bloomington stock center) each of which is associated with an image of the neuronal expression pattern shown at http://flweb.janelia.org/cgi-bin/flew.cgi. In addition we used the insertion site stocks, w;attP2 and w;attP2;attP40(Pfeiffer et al., 2008; 2010), ppk1.9-GAL4 (Ainsley et al., 2003), 19-12-GAL4, nompC (Zhang et al., 2013) and iav-GAL4 (Kwon et al., 2010). We used the progeny larvae from the insertion site stock, w;;attp2, crossed to the appropriate effector (UAS-TNT-e (II)) for characterizing the w;; attP2 were selected because they have the same genetic background as the GAL4 tested in the screen. We used the following effector stocks: UAS-TNT-e (Sweeney et al., 1995) and pJFRC12-10XUAS-IVSmyr::GFP (Bloomington stock number: 32197).

### Larva dissection and immunocytochemistry

To analyze the expression pattern of the GAL4 lines, we crossed the lines to pJFRC12-10XUAS-IVSmyr::GFP (Bloomington stock number: 32197; [23]). The progeny larvae were placed in a phosphate buffered saline (PBS; pH 7.4) and fixed with 4.0% paraformaldehyde for 1-2 hr at room temperature, and then rinsed several times in PBS with 1% Triton X-100 (PBS-TX). Tissues when then mounted on poly-L-lysine(Sigma-Aldrich) coated coverslips and then transferred to a coverslip staining JAR (Electron Microscopy Sciences) with blocking solution, 3% normal donkey serum in PBS-TX for 1 hr. Primary antibodies were used at a concentration of 1:1000 for rabbit anti-GFP (Invitrogen) and 1:50 for mouse antineuroglian (Developmental Studies Hybridoma Bank) and 1:50 for anti-N-cadherin (Developmental Studies Hybridoma Bank) and incubated for 2 days at 4°C. Tissues when rinsed multiple times in PBS-TX and then incubated for 2 days. with secondary antibodies: anti-mouse IgG Alexa Fluor 568 Donkey (diluted 1:500; Invitrogen), Alexa Fluor 647 Donkey anti-rat IgG (1:500, Jackson ImmunoResearch) and fluorescein FITC conjugated Donkey anti-rabbit (diluted 1:500; Jackson ImmunoResearch). After incubation, the tissue was rinsed for several hours in PBSTX, and dehydrated through a graded ethanol series, cleared in xylene and mounted in DPX (Sigma) Images were obtained with 40x oil immersion objective (NA 1.3) on a Zeiss 510 Confocal microscope. Images of each nervous system were assembled from a 2xarray of tiled stacks, with each stack scanned as an 8 bit image with a resolution of 512x512 and a Z-step of 2 μm. Images were processed using Fiji (http://fiji.sc/)

### Behavioral apparatus

The apparatus was described previously (Jovanic et al., 2016; Ohyama et al., 2013). Briefly, the apparatus comprises a video camera (DALSA Falcon 4M30 camera) for monitoring larvae, a ring light illuminator (Cree C503B-RCS-CW0Z0AA1 at 624 nm in the red), a computer (see (Ohyama et al., 2013) for details); available upon request are the bill of materials, schematic diagrams and PCB CAM files for the assembly of the apparatus) and a hardware modules for controlling air-puff, controlled through multi worm tracker (MWT) software (http://sourceforge.net/projects/mwt) (Swierczek et al., 2011), as described in (Ohyama et al., 2013). Air-puff is delivered as described previously (Ohyama et al., 2013). Briefly it is applied to a 25625 cm2 arena at a pressure of 1.1 MPa through a 3D-printed flare nozzle placed above the arena (with a 16 cm 6 0.17 cm opening) connected through a tubing system to plant supplied compressed air (0.5 MPa converted to a maximum of 1.4 MPa using a Maxpro Technologies DLA 5-1 air amplifier, standard quality for medical air with dewpoint of 210uC at 90 psig; relative humidity at 25uC and 32uC, ca. 1.2% and 0.9%, respectively). The strength of the airflow is controlled through a regulator downstream from the air amplifier and turned on and off with a solenoid valve (Parker Skinner 71215SN2GN00). Air-flow rates at 9 different positions in the arena were measure with a hot-wire anemometer to ensure even coverage of the arena (Extech Model 407119A and Accusense model UAS1000 by DegreeC). The air-current relay is triggered through TTL pulses delivered by a Measurement Computing PCI-CTR05 5-channel, counter/timer board at the direction of the MWT. The onset and durations of the stimulus is also controlled through the MWT.

### Behavioral Experiments

Embryos were collected for 8–16 hours at 25°C with 65% humidity. Larvae were raised at 25°C with normal cornmeal food. Foraging 3 instar larvae were used (larvae reared 72-84 hours or for 3 days at 25°C).

Before experiments, larvae were separated from food using 10% sucrose, scooped with a paint brush into a sieve and washed with water (as described previously). This is because sucrose is denser than water, and larvae quickly float up in sucrose making scooping them out from food a lot faster and easier. This method is especially useful for experiments with large number of animals. We have controlled for the effect and have seen no difference in the behavior between larvae scooped with sucrose and larvae scooped directly from the food plate with a forceps.

The larvae were dried and spread on the agar starting from the center of the arena. The substrate for behavioral experiments was a 3% Bacto agar gel in a 25625 cm2 square plastic dishes. Larvae were washed with water at room temperature, the dishes were kept at room temperature and the temperature on the rig inside the enclosure was set to 25°C.

The humidity in the room is monitored and held at 58%, with humidifiers (Humidifirst Mist Pac-5 Ultrasonic Humidifier).

We tested approximately 50–100 larvae at once in the behavioral assays. The temperature of the entire rig was kept at 25 °C. In the assay, the larvae were left to crawl freely on an agar plate for 44 seconds prior the stimulus delivery. The air-puff was delivered at the 45^th^ second and applied for 38 seconds. After a period of recovery of 22 seconds when 10 air-puff pulses, 2 second each, were delivered (with a 8 second separation interval). WE computed behavioral and transitional probabilities for different time-windows during stimulation. In this study, for the data from the behavioral screen, we show behavioral probabilities for the first seconds after stimulus onset and the transition probabilities for the first 3 seconds after stimulus onset.

The MWT software 64 (http://sourceforge.net/projects/mwt) was used to record all behavioral responses.fcast

### Screen design

We screened 567 GAL4 lines from the Rubin GAL4 collection ((Jenett et al., 2012; Pfeiffer et al., 2008) were we silenced small subsets of neurons and individual neurons using tetanus toxin. We selected these lines from the entire collection for sparse expression in the brain and ventral nerve cord of the larval CNS as well as expression in the sensory neurons (images of the larval CNS are available at http://www.janelia.org/gal4-gen1). Out of the 567 lines tested, there were neuronal lines that were not part of the collection: we added 19-12 GAL4 and nompC-GAL4 for sensory neurons and OK107GAL4 for the mushroom body. We screened each GAL4 line in the air-puff assay described above.

### Behavioral analysis

#### Larva tracking

Larvae were tracked in real-time using the MWT software (Swierczek et al., 2011). We rejected objects that were tracked for less than 5 seconds or moved less than one body length of the larva. For each larva MWT returns a contour, spine and center of mass as a function of time. From the MWT tracking data we computed the key parameters of larval motion, using specific choreography (part of the MWT software package) variables (Ohyama et al., 2013).

#### Behavioral detection

From the tracking data, we detected and quantified the behaviors: head-retractions, head-casts (Cast), backwards crawls (Back-ups), stopping (Stop) and peristaltic crawling strides (Crawl) using a behavior classification that allows discriminations between all the different actions. Behavior classification is performed using supervised learning based on human tagging of larval videos i.e. (Kabra et al., 2012). It is performed on a very limited set of features exhibiting low variance under the mechanical deformations induce by the larval dynamics. It consists on a 3-layer procedure. The first layer relies on Random Forest (Breiman, 2001) to identify if one of the listed behavior is being performed and output a Boolean variable. The second layer collects all states and check for inconsistencies (e.g. a larva crawling and head-casting at the same time). The third layer used Random Forest again to perform the final behavior assessment (Bishop, 2006)

#### Tagging data

Data tagging was performed using custom, simple GUI in Matlab. A first version provided the path of the larva, the contour and spine. The head, tail (non differeniated) and neck points were apparent. The evolution of 2 features could be plotted to help tag some actions. Tagging was limited to whether an action was ongoing or not.

A second GUI (the 2 versions of it) with similar design displayed the state assigned by the previous layer of classification, the differentiated positions of the head and tail as well. One version was done with a local zoom on the larva to provide 2 simultaneous view, one more focused on global dynamics and one more local.

#### Principle and method used to action detection

The analysis is done in sequential layer using the following simple architecture:

- Loading and Cleaning input files
- Generate features time series
- Identify Head and Tail
- Generate first estimation of the actions
- Regularize these actions using imposed hierarchy and custom rules
- Generate second estimate of the actions
- Regularize singular events
- Action specific corrections

#### Loading and cleaning input files

Inputs of the analysis pipeline are made of the time series of the contours and the spine of individual larva as produced by the Choreography scheme (Ohyama et al., 2013). Contours are dynamically generated live on subsets of larva from very different size and shape and thus have non-constant size. The average length is 114±23 points measures on a sample of 50 000 larvae. Spines, which are the skeleton of the larva, were generated offline and were all made of 11 points. Input files were then cleaned of the 5 times first points and last 10 time points which bear usually several anomalies (at the screen scale). We removed individual larva experiments that had not at least 250 points (lasted ~ 20s) and whose trajectories had a convex hull of surface inferior to 1mm^2^ (nearly did not move at all).

#### Generate features

Complete details of all features computed are shown in the (https://github.com/DecBayComp/Pipelineactionanalysist5pasteurjanelia) Here, we list key features that provided selectivity in classification.

The center of the larva was defined as 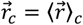 with 〈 〉_*c*_ the averaging along the contour.

The “necks” of the larva were defined as 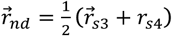, 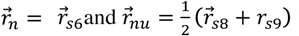 where the index *nd* stands for neck-down, *n* for neck, *nu* for neck-up and with 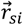 is the ith point of the spine.

We defined 4 segments in the larva, *S*_*e*1_ joining the head to the neck-up, *S*_*e*2_ joining the center to the neck up, *S*_*e*3_ joining the neck-down to the center and *S*_*e*4_ the segment joining the tail to the neck-down.

We defined 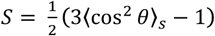 with 〈 〉_*s*_ the averaging along the spine curve, cos *θ* the scalar product between normalized vector associated to a segment of the spine and the direction of the larva body define as the normalized vector joining the lower neck the mid point of the spine. S takes value between −0.5 and 1.

We defined 4 main angles as *θ*_1–4_ as respectively, *θ*_1_ the angles between *S*_*e*1_ and *S*_*e*3_, *θ*_2_ the angles between *S*_*e*4_, *S*_*e*2_, *θ*_3_ the angles between *S*_*e*1_ and *S*_*e*2_, *θ*_4_ the angles between *S*_*e*3_ and *S*_*e*4_.

We defined 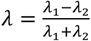, with *λ_i_* the eigenvalues of the matrix 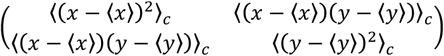 with *λ*_1_ ≥ *λ*_2_. *λ* characterizes the shape of the larva and takes value between 0 and 1.

Various velocities or acceleration were evaluated both directly and then smoothed or by convolution with derivatives of Gaussian with different standard deviations. We evaluated the effective agitation velocity as 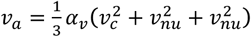 with *α_v_* = 10 for this screen analysis. We also evaluated the motion velocity and acceleration of the head, tail and center of mass of the larva.

We defined a set of linked variables defining direction of motion

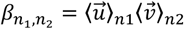

with 〈 〉*_n_i__*. the temporal Gaussian averaging over *n_i_* points, 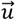 the unit vector going from tail to neck down and 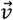 the unit vector in the direction of motion (center of mass).

We defined the various scale effective distance moved

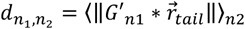

with *G′* the derivatives over *n*_1_ point, * the convolution, || || the norm, and 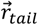 the position of the tail

We defined the true and modified length of the larva as 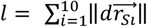 and 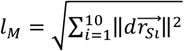 with 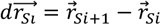.

We defined the effective rotation energies

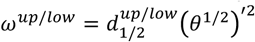

with 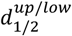 the distance between head/tail and the center of the larva, 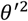 the squared derivative of the angle.

We defined the auxiliary direction variables as 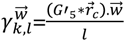 with 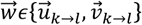, {*k* → *l* }*𝜖*{neck → head, neck → tail, tail → head } and 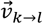 the vector perpendicular to 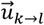.

Features were then smoothed at various time scales, derived, log-derived, and squared after derivation. Furthermore, minimal, maximal and cumulative values over time points window of various size were evaluated.

Some features were also dynamically normalized with respect to the length of the larva. Motion velocities were normalized by 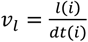. The surface of the contour was split into two normalized variables, one normalized by the surface of convex hull of the contour the other by the surface of the circle of radius 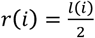. The direct distance from head to tail, the distance from the head to the neck and the neck to the tail were normalized by the length of the larva *l*(*i*).

Finally, last layer classifications used global variables averaged over the full temporal duration of the larva dynamics: 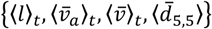 where only the length of the larva is not normalized.

#### Identify of Head and Tail

In order to properly sample the large space of possible actions and to explore differences between individual larva, experiments were performed on large sets of larva. To reduce probability of contact between larva that would induced unwanted actions, agar plates were made large enough to have large portion of “free” motion during times or recording. In order to match all these conditions, the optical magnification was chosen to be low providing limited features on the larva. Thus, neither the mouth hook nor the difference of curvature between head and tail could not be reliably detected. Hence, neither individual images nor contour shapes could be reliably used to identify head and tail.

Head and tail are the terminal point of the spine; hence both head and tail can be in position 1 or 11. We relied on a Hidden Markov model (HMM) to detect head and tail from the larva dynamics. To reduce the dynamics to a one time step HMM, 4 hidden states were introduced corresponding to the 4 possible transitions between 2 time steps: T_1_: (Head 1 → 1, Tail 11 → 11), T_2_: (Head 1 → 11,Tail 11 → 1), T_3_: (Head 11 → 11,Tail 1 → 1), and T_4_: (Head 11 → 1,Tail 1 → 11).

The likelihood model was defined to reinforce global characteristics of head and tail dynamics when recorded in high resolution. The likelihood at the time point i reads:

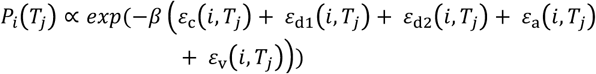

with

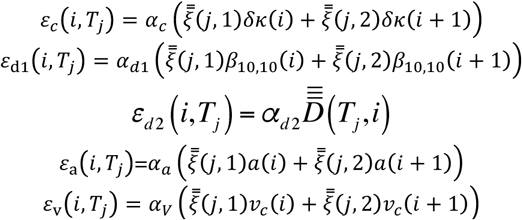

with the matrix 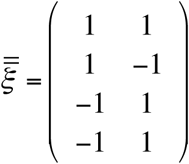, 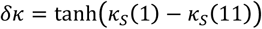 with *K* is in radians and defined in Schulze et al., 2015,

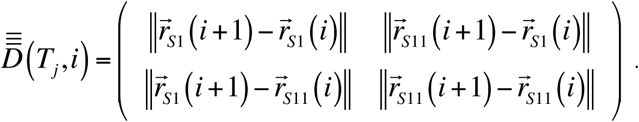

In order to regularize the contour of the larva and evaluate the curvature at the Head/Tail points we projected and reconstituted the larva contour onto a Fourier basis. This procedure was directly inspired by a step of the closed-loop analysis code found in Schulze et al., 2015. We limited the decomposition and recomposition to the order 7 in coefficients. Curvatures were measured using the reconstituted (smooth) larva contour.

Parameters were set to *α* = (1,0.2,1,0.5,0.5), *β* = 0.01, 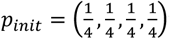 with a transition matrix 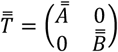 with 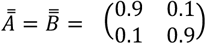. The transition matrix was purposefully set to much higher transition values than temporally observed. It allowed possible changes following complex maneuvers. Note that the same value was used at the screen scale. Another analysis could be performed by adapting the transition matrix to the shape and dynamics of the larvae.

The head and tail position were derived from the Viberti path (path of hidden states maximizing the likelihood). As for all HMM process the longer the trajectory the better the results.

The main source of error stemmed from the transition from ball form to a moving dynamic. When the larva is curled the spine is constantly moving due to the lack of high curvature area to stabilize it. If the larva performs a ball shape moves for a short time and then performs a ball again, there are limited constrain to define head and tails.

Notably, only for the crawling backward is the head and tail definition effectively used.

#### The multilayer architecture

We point out here the reason for the multiple layers of classification. Behavior or action tagging in larva by human observers is not fully reliable. Different people tend to see different dynamics when looking at the contour and time series dynamics. Finally, in many cases the low resolution of the contour makes the choice of dynamics unsure. Hence, first layers of classification were performed on limited larva exhibiting clear dynamics and any unsure actions were not tagged. These actions were then used to classify actions at the full screen scale. This allowed future tagging to be performed with the first layers state being displayed. This process helped to unify description of the dynamics and provided clearer elements to judge the action dynamics.

#### Generate first estimation of the actions: First layer classification

In order to reduce the effect of misclassification of the head and tail, all classification except the ones explicitly for crawling backward were done without requiring their detections.

Classifications were performed using Random Forest on the small samples of tagged larva. Selected sets of features were provided in relation to the action to be tagged. Action detection was performed independently for each action with results being at each time point the action being on or off. The 5 actions and additional actions were detected.

Additional postural states were also detected. These postural states simply describe non-dynamic the shape of the contour. They were divided into 4 non-overlapping state: straight, cast, curl and ball. An additional fifth state straight and light cast was added to improve backward motion detection especially when associated in sequences involving cast and back-up.

Crawling backward was defined as the combination of the straight and light bent (cast) postural state, the displacement status and the feature axis_direction_25_10 taking values inferior to the threshold −0.8.

#### Regularized actions using imposed hierarchy and custom rules

The first layer of tagging was performed on limited set of larvae. Hence, we left at this stage of the code, the possibility of having a state with no action classification. Furthermore, each action was defined independently of the other. This no action set was defined hierarchically transferring

- Any action happening at the same time as stop was set to no-action
- Any action happening at the same time as roll was set to no-action
- Any action happening at the same time as back-up was set to no-action
- Any action happening at the same time as cast was set to no-action
- Any action happening at the same time as crawl was set to no-action

Similarly, postural states were regularized hierarchically, using the probability associated to the Random Forest as defining criteria. The postures were regularized sequentially in the following order ball → curl → cast → straight.

#### Generate second estimate of the actions: Second layer classification

The second layer classification was based on Random Forest. All actions were trained at the same time with an output being of the 6 actions. All actions were tagged and the no-action states were removed. Finally, to the time series of features computed on the larva contour and spine, the actions classification from the previous layer and their smooth probabilities were added as an input of the random forest. Auxiliary actions and states were not re-trained on this layer.

### Regularize singular events

The nature of the larva dynamics and the scale (temporal and spatial) at which the behavior is analyzed prevent certain transition to be possible. We thus regularized two patterns: iji and ijk where {i,j,k} are one of the possible actions, and where these actions are separated by 1 time point. The iji pattern was systematically corrected to iii and the ijk pattern to ikk.

### Action specific corrections

Action specific corrections are based on evaluating parameters during the action as defined in the previous layers.

### Head-retraction

New features were generated to ensure a better discrimination when the action head retraction was detected. These features were associated to the length of the larva and to the average, minimum and maximum values taken during the detect head retraction action. A random forest was the train to differentiate between head retraction and other quick motions that could be taken as head retraction.

### Stop

Stops lasting less than 2 time points were transformed into the closest action they could be performing.

### Back-ups

Back-ups lasting less than 2 time points were transformed. This correction was implemented to lift the ambiguity of nearly stopped larva where small back mostly consequences of the resolution of the tracker were generated.

### Action continuum vs discrete actions

Larva behavior was analyzed at the screen scale under the discrete actions hypothesis. This hypothesis is supported by natural clustering on simple analytical basis. In Supplementary figure 1, we showed that using a subset of parameters and a t-sne embedding (van den Maaten and Hinton, 2008), natural clusters appear corresponding to 5 main actions performed by the larva. Coloring was performed using the pipeline. The 6 parameters used 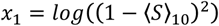, 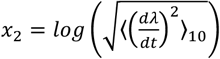, 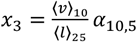, 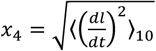, 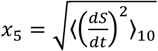, 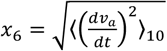 with 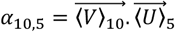, where 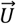 is the unit vector going from the tail to the mid point of the spine and 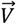 is the unit vector in the direction of motion of the mid-point of the spine (the direction of motion). Each point in the plots is the average value of the parameters in one experiment when the larva performs on the behavior.

Here, we show multiple embedding with various number of larva included in the reference. We also show examples with few other lines. Hence, average in time these parameters allow clustering of the dynamics.

### Action Definition

#### Stop

A larva is stopped if it does not show any detectable muscular activity. A non-crawling larva is stopped if it does not dynamically bend. A stopped larva can be in a bent posture.

#### Head retraction

A larva is Head-retracting if it is dynamically (and most of the time more quickly than the rest of the larva dynamics) diminishing its length beyond peristaltic wave amplitude. Head retraction is not associated to a backward movement.

#### Cast

Any dynamics of bending in the larva whether it is displacing (turning movements) or not displacing (head or tail casting movements). Here, displacing means the center of mass or the center of the larva (not necessarily the main point) is moving.

#### Crawl

Larva displacement in the direction going from the tail to the head.

#### Back-up

Larva displacement in the direction going from the head to the tail.

#### Rolling

A larva is rolling it is moving laterally while being bent. Rolling is not present in this study (Roll is a type of nociceptive response, that doesn’t normally occur in response to air-puffs.

### Choice in Data used for training

The Behavioral larva database is made of times series of larva contour and larva spine. All are untagged. Probability of actions in drosophila larva is extremely heterogeneous and some actions can only be seen following specific stimuli patterns. For example a sample of nearly 14000 larvae taken in the reference (w;;attP2) exhibit the proportion of events follow the following raw statistics: 50% crawling, 37% casting, 9% stopped, 1% head retracting, 3% crawling backwards (Back-ups). Hence, in order to properly sample a large variety of events tagging was done selectively with a focus on pre and post stimuli (the main at 45 s but also the series of 10 stimuli beginning at 95s). First set of data also included larva from an activation screen (unpublished data) and similarly we focused tagging around the activation time. Tagging for the last layer was exclusively done on larva from the current screen. Initial tagging was performed on “hits” and reference lines described in (Jovanic et al., 2016; Ohyama et al., 2013) and the above said activation screen (upublished). Then, random sampling of the screens was performed.

### Temporal Anomaly

Note that a dependency exists in the MWT system between the number of larvae actively tracked and the time between frames. The average time between frames is dt= 8.2 10^−2^± 1.3 10^−2^ s with minimal value of dt_min_=2.8 10^−2^ s and maximal value of. dt_max_ =2.9 s. These values were evaluated on a sample of 50000 larvae and total corresponding recording time of T=56000 s randomly sampled from the full experimental database. Local variability of dt may degrade the quality of action assessment.

### Anomalous locomotor defect phenotypes

We defined anomalous phenotypes at the population scale. Nonstimulated larva moving freely on agar plates exhibits two actions: forward crawling with an average time of ~10s and casting with an average time of ˜1s. These two dynamics follow roughly a Poisson dynamics. Hence at any time, at the population scale crawling is the dominant behavior (the main behavior happening) and the second dominant is cast. We removed experimental sets if this hierarchy was violated prior to the stimuli at 45s.

### Evolution of action definition

In this paper action description slightly deviated from the ones in (Jovanic et al., 2016; Ohyama et al., 2013) while remaining compatible. We provide here, the transition between the two.

### Head retraction vs Hunch

In previous studies (Jovanic et al., 2016; Ohyama et al., 2013), “hunch” was defined as the strong length decrease of the larva with the tresholds as describe in (Ohyama et al., 2013). “Head-retraction”, as defined in the present study, are all events of larval shortening regardless of their magnitude. If we apply a similar threshold (effective length change dl=0.8) used for previous “hunch” detection with the present machine-learning behavioral detection method we obtain previously defined hunches.

### Cast vs Bend

Similarly, previously (Jovanic et al., 2016) “bends” were described as events with head-tail angles of 27° or more (Ohyama et al., 2013). If we apply a similar threshold to the “cast” events detected with the present machine-learning algorithms, we obtain the previously defined “bends” (effective angle change, S=0.81).

## Quantification and statistical analysis

### Ethograms

Ethograms were designed to display the temporal evolution of the individual larva behavior with time. Color code is associated to the action being performed. Stimuli are displayed in magenta thick lines.

### Behavior probability

The temporal evolution of behavior was computed as the ratio of the number of larva performing the action over the total number of active larva in time windows of duration 2dt with dt=100ms. Noticeable evolution of noise in the curves may be the result of varying number of active larvae in the rig.

### Dominant Behavior

The temporal evolution of dominant and second dominant behavior was computed as the most probable and second most probable behavior in same time window as behavior probability.

### Transition analysis

Transition analysis simply consisted of counting during a predefined time window all the transition between behaviors. These transitions are represented in matrices where initial states are rows and arrival states are columns. Normalization was performed on rows. The transition probabilities starting from a given behavior for control and experimental populations were statistically compared by the Fisher exact test. We say that any p-value less than 0.05 (uncorrected for multiple comparisons) is significant.

### Sequence analysis

Sequences were measured as the list of different actions performed after the stimulus. We limited the number of actions to the first four actions performed after stimulus onset. We defined dt_blur_= 500ms as the time of uncertainty if the stimulus has been perceived. If the larva performs the same actions as it was before the stimulus for longer than dt_blur_ it is considered to be the response to the stimuli. We included in this analysis all the actions: head-retractions, casts, back-ups, stops as well as the baseline behavior crawl.

We used a chi-squared statistical test to compare GAL4 lines with control (w;;attP2). We say that any p-value less than 0.05 (uncorrected for multiple comparisons) is significant.

### Behavioral probability analysis (time windows)

Probability analysis was performed on averaging on various time windows, 1s, 5s and 15s, the fraction of time that the larvae spent performing an action. This analysis was performed after the start and the end of the stimuli.

We used a chi-squared statistical test to compare GAL4 lines with control (w;;attP2). We say that any p-value less than 0.05 (uncorrected for multiple comparisons) is significant.

### Cumulative analysis

Cumulative analysis was performed by counting the number of larva performing at least once an action in various time widows of duration 1s, 5s, and 15s. This analysis was performed after the start and the end of the stimuli.

### Larva length analysis

Length analysis was performed during head retraction event. We evaluated the maximal and minimal length (both *l* and *l_M_*) during the head-retraction and measured the relative variation 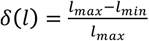 and 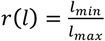.

### Amplitude during cast

The amplitude of cast was analyzed during cast event after the stimulus. We estimated the average value of the minimal S value during the cast.

### Hit detection

Behavioral probability and sequence probability hits were considered the 10% of animals with highest and lowest behavioral probabilities that were significantly different from the control (w;;attP2). We excluded from the hits all the lines that have a locomotor defect (see Supplementary spreadsheet_locomotor defect), all the lines that didn’t have at least 40 larvae from at least 2 valid repetitions. The behavioral probability hits when then divided into 3 categories as described in Figure 2. Depending on whether the lines shows an increase or decrease in behavioral probabilities or both the hits were categorized in less response, more response and decision or competitive interaction hits

Out of all the lines that had significantly different transition probabilities (for at least one type of transitions) for the purpose of this study, we considered as hits the lines in which silencing affected only one type of transition or reversal (reversal-transition back onto the previous action). We also investigated hits where both head-retraction to casts and head-retraction to back-ups were affect upon silencing.

## Acknowledgments

We thank C.Montell (UC Santa Barbara), YN Jan (UC San Francisco) for fly stocks. We thank Fly core (especially Monti Mercer) at JRC for fly crosses and Rebecca Arruda and Tam Dang for help with the behavioral experiments.

## Supplementary tables and spreadsheets

Supplementary spreadsheet_locomotor defects (abnormal behavior prior to stimulus onset) Supplementary spreadsheet_behavioral probabilities

Supplementary spreadsheet_transition probabilities - Transition probabilities screen scale. For each transition probability a p-value for comparison with the corresponding transition probability in the control (w;; attp2) are shown. P-values <0.05 are shown in red

**Supplementary table 1.** Sensory line hits. Behavioral probabilities and p-values for comparison of each of the lines with the control

**Supplementary table 2.** CNS line hits. Behavioral probabilities and p-values for comparison of each of the lines with the control

**Supplementary table 3.** Transition probabilities hits. For each transition probability a p-value for comparison with the corresponding transition probability in the control (w;; attp2) are shown. P-values <0.05 for transition are shown in red, reversals are shown in green. The dark yellow labels the transition probabilities that are higher than the control

**Supplementary table 4.** The 10 most represented 4 action sequence motifs in the w;;attp2-TNT control that occur in response to an air-puff. The four actions: head-retraction, cast, back-up, stop and the baseline behavior crawl are shown

---

**Supplementary figure 1.**
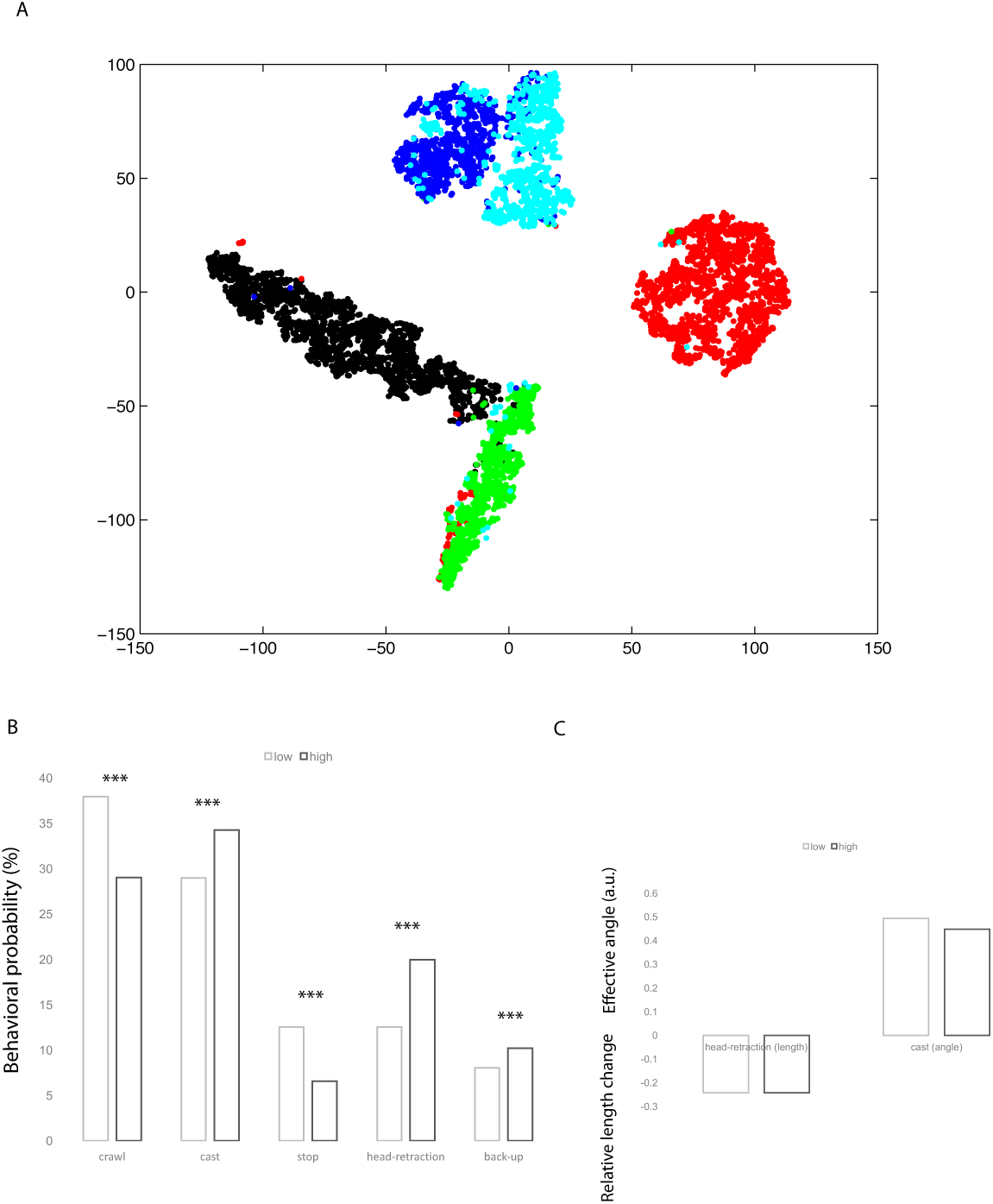
**A.** Natural clustering. Discrete actions emerge naturally as separate clusters in a simple embedding space derived from few features. X and Y axis are arbitrary variables generated during the t-sne (van den Maaten and Hinton, 2008) computation. **B.** Behavioral probabilities (1^st^ second) at high (6 m/s) and low (3 m/s) intensity of air-puff. P-values are all p<0.001 (***) **C.** Amplitude of casting and head-retractions and high and low intensity of air-puff

**Supplementary figure 2.**
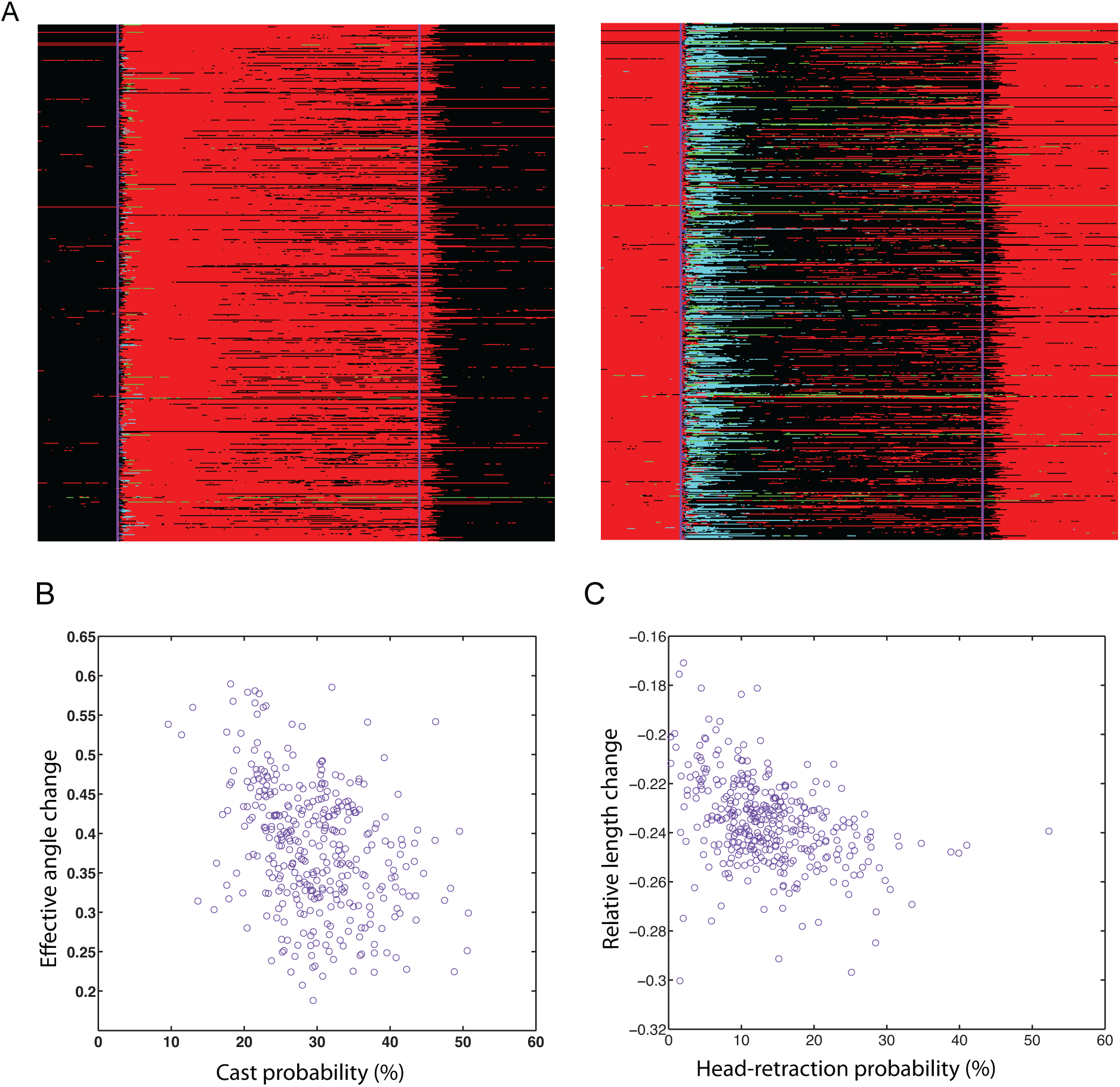
**A.** Ethogram of the behaviors of all the lines from the screen before, during (38s) and after stimulation-dominant behavior is shown in the ethogram on the left and second dominant behavior in the ethogram on the right. Each line represents a neuronal line. **B.** Scatterplots of Amplitude of Casting (effective angle) and Cast probability **C.** Scatterplots of Amplitude of head-retractions (Relative length change) and head-retraction probability. **B-C.** Each dot represents the average value for a neuronal line.

**Supplementary figure 3.**
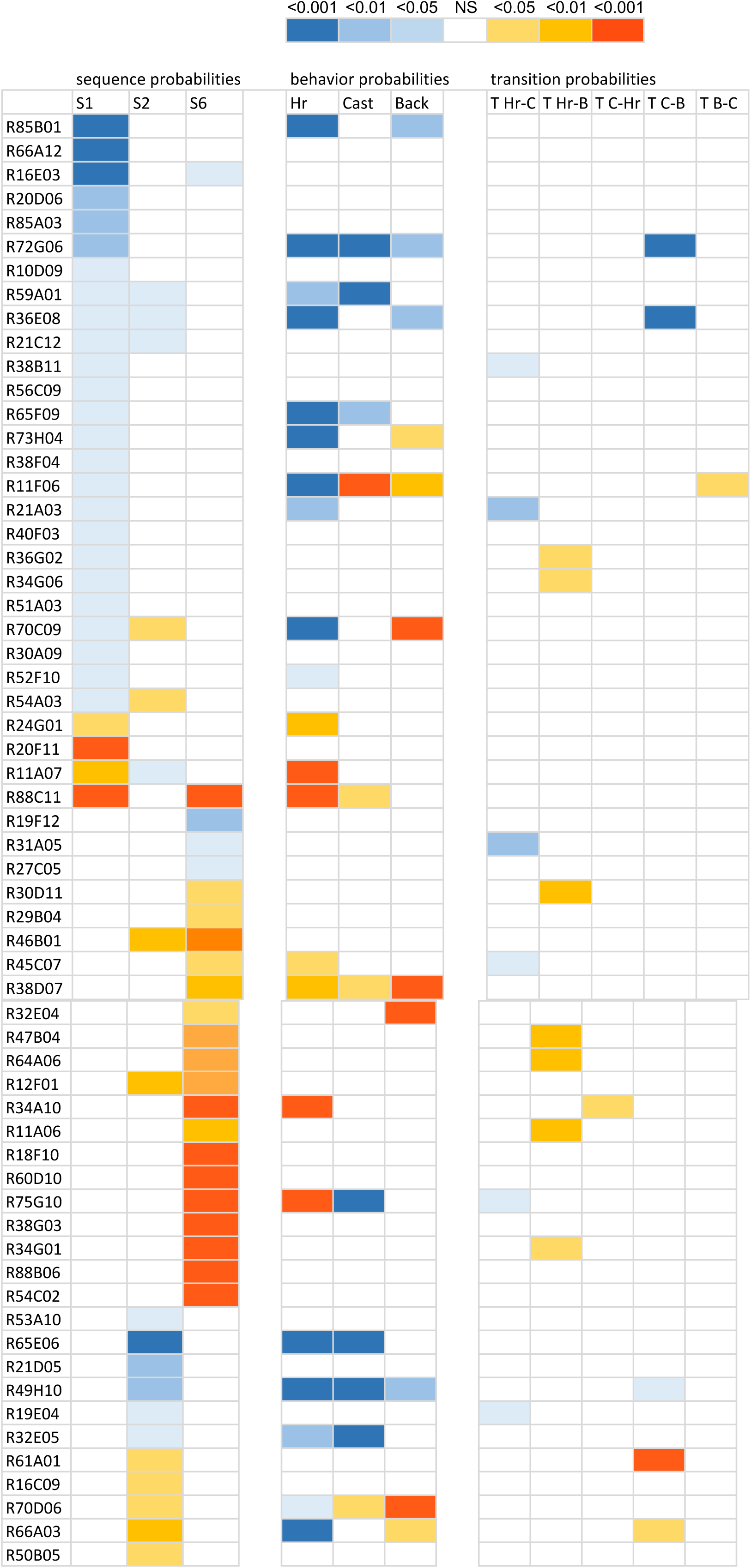
S1-S3 sequence motifs probabilities. S1: Head-retraction – Cast - Back-up - Head-retraction, S2: Cast - Back-up – Cast - Back-up, S3: Head-retraction – Back-up – Cast- Back-up, Hr, cast and Back – behavioral probabilities of head-retraction, Casting and Backing-up respectively. T transition probabilities between pairs of actions as indicated (Hr-head-retraction, Bback- up and C - cast 61 lines that have sequence motifs probabilities S1-S3 significantly higher or lower compared to the control are shown. 34 of these lines had also behavioral probabilities and/or transition probabilities involving head-retraction, cast and back-up, affected (significantly different than the control). 25 lines had decreased Head-Retraction-Cast-Back-up-Cast probabilities out of which five had also the probability Head-retraction-Back-up-Cast-Back-up or the probability of Cast-back-up-Cast- Back-up affected. 4 have an increased Head-Retraction-Cast-Back-up-Cast motifs, out of which 1 had the Cast-back-up-Cast- Back-up probability decreased and one had a Head-retraction-Back-up-Cast-Back-up increased. 19 lines had the had the probability of Head-retraction-Back-up-Cast-Back-up (out of which 2 had the Cast-back-up-Cast-back-up increased) motifs increased while 3 had that same probability decreased. The probability of the Cast-Back-up-Cast-Back-up-Cast was increased in 5 lines and decreased in 6 lines.

